# The LRRK2 kinase substrates Rab8a and Rab10 contribute complementary but distinct disease-relevant phenotypes in human neurons

**DOI:** 10.1101/2023.04.30.538317

**Authors:** Adamantios Mamais, Anwesha Sanyal, Austin Fajfer, Catherine G. Zykoski, Michael Guldin, Alexis Riley-DiPaolo, Nitya Subrahmanian, Whitney Gibbs, Steven Lin, Matthew J. LaVoie

## Abstract

Mutations in the LRRK2 gene cause familial Parkinson’s disease presenting with pleomorphic neuropathology that can involve α-synuclein or tau accumulation. LRRK2 mutations are thought to converge toward a pathogenic increase in LRRK2 kinase activity. A subset of small Rab GTPases have been identified as LRRK2 substrates, with LRRK2-dependent phosphorylation resulting in Rab inactivation. We used CRISPR/Cas9 genome editing to generate a novel series of isogenic iPSC lines deficient in the two most well validated LRRK2 substrates, Rab8a and Rab10, from two independent, deeply phenotyped healthy control lines. Thorough characterization of NGN2-induced neurons revealed divergent effects of Rab8a and Rab10 deficiency on lysosomal pH, LAMP1 association with Golgi, α-synuclein insolubility and tau phosphorylation, while parallel effects on lysosomal numbers and Golgi clustering were observed. Our data demonstrate largely antagonistic effects of genetic Rab8a or Rab10 inactivation which provide discrete insight into the pathologic features of their biochemical inactivation by pathogenic LRRK2 mutation.

**Highlights:**

- Rab8a and Rab10 deficiency induce lysosomal and Golgi defects
- Rab8a and Rab10 deficiency induce opposing effects on lysosomal pH
- Rab8a KO and Rab10 KO neurons show divergent effects on synuclein and tau proteostasis
- Inactivation of different Rab GTPases can induce distinct disease-relevant phenotypes

## Introduction

Missense mutations in the LRRK2 gene are the most common cause for autosomal dominant Parkinson’s disease (PD), with this locus further linked to idiopathic PD risk (Barrett et al., 2008; Bonifati, 2006; Paisán-RuÍz et al., 2004). LRRK2 is a large, 2,527 amino acid protein with functional GTPase and kinase domains where the PD-linked mutations largely cluster (Kluss et al., 2019). Upon neuropathologic examination, LRRK2-PD can manifest with pure nigral degeneration (rare) or classic Lewy body pathology typical of idiopathic PD, but surprisingly a majority of cases show deposition of tau into neurofibrillary tangles (Henderson et al., 2019; Herbst et al., 2022; Ujiie et al., 2012; Zimprich et al., 2004). How LRRK2 pathobiology might provoke distinct and independent neuropathological features is one of the greatest unanswered questions in the field.

A large subset of Rab GTPases are amongst the most well-established LRRK2 substrates. LRRK2-dependent phosphorylation occurs at a highly conserved threonine residue in their GTPase switch domain which results in a net inhibition of Rab activity (Pfeffer, 2018; Steger et al., 2016). Rab GTPases orchestrate the spatiotemporal regulation of intracellular membrane and protein trafficking across the cell and can confer membrane identity, organelle morphology and organelle protein content (Pfeffer, 2018; Stenmark, 2009). Rab mutations have been linked to various diseases including cancer, Charcot-Marie-Tooth disease type 2B, and Warburg Micro syndrome (Banworth and Li, 2017). Thus, their mis-regulation by mutant LRRK2 activity is well-positioned to broadly impact cellular proteostasis that may contribute to neurodegenerative disease.

Rab GTPases show selectivity for different cellular compartments in endocytic and recycling pathways conferring divergent functions (Pfeffer, 2018). Amongst the ∼14 Rab proteins thought to be phosphorylated by LRRK2, Rab8a and Rab10 are the most well-validated to date. Rab8a has been linked to neurodegeneration via its interactions with the PD associated proteins α-synuclein and PINK1 (Vieweg et al., 2020; Yin et al., 2014), in addition to phosphorylation by LRRK2 (McFarland et al., 2018; Vieweg et al., 2020). While our understanding of their function is incomplete, both Rab8a and Rab10 have been implicated in the cellular response to lysosomal injury or stress (Bonet-Ponce et al., 2020; Eguchi et al., 2018; Mamais et al., 2021). Interestingly, Rab8a and Rab10 have opposing roles in normal ciliogenesis (Dhekne et al., 2018), but such antagonistic properties have not been explored in other contexts. Taken together, these data endorse the parallel consideration of Rab8a and Rab10 biology as performed here and highlight their potential importance in LRRK2-dependent disease.

Here, we used CRISPR/Cas9 genome editing to ablate Rab8a or Rab10 expression in iPSCs derived from deeply phenotyped (clinically and genetically) wild-type healthy control human subjects (Bennett et al., 2018; Lagomarsino et al., 2021). Multiple isogenic clones from two independent control lines were differentiated into cortical neurons by forced Neurogenin-2 (NGN2) expression to produce induced neurons (iNs). We report a significant decrease in lysosomal number across both Rab8a KO and Rab10 KO neurons compared to their isogenic controls. However, Rab8a KO neurons exhibited mild lysosome acidification whereas Rab10 KO lysosomes were mildly alkaline. Altered glycosylation of the lysosomal protein LAMP1 was noted in both Rab8a KO and Rab10 KO models, prompting an investigation into Golgi integrity. Morphological analysis of cis and trans Golgi networks revealed a compressed, or clustered, Golgi distribution in Rab8a and Rab10 KO iPSCs and a concomitant increase in LAMP1 retention in the trans Golgi network in Rab8a KO cells.

LRRK2-PD most commonly manifests with either classic Lewy body inclusions comprised of α-synuclein, or neurofibrillary tangles composed of hyper-phosphorylated tau. Given that these two proteins are selectively expressed in neurons and their deposition in disease occurs across multiple neuronal subtypes, we explored the consequence of Rab8a or Rab10 deficiency in iNs. Consistent with other opposing phenotypes linked to these two LRRK2 substrates, we found features of α-synuclein homeostasis altered in Rab8a but not Rab10 KO neurons, with selective effects on Tau in Rab10 but not Rab8a KO neurons. These data not only identify distinct cellular pathways that likely contribute to the unique pleiomorphic pathology associated with LRRK2-PD, but may also address why co-morbid α-synuclein and tau pathology is not reported (Henderson et al., 2019; Herbst et al., 2022; Ujiie et al., 2012; Zimprich et al., 2004).

## Results

### Divergent lysosomal alterations in Rab8a KO and Rab10 KO human neurons

To establish neuronal models of Rab8a and Rab10 deficiency we used CRISPR/Cas9 gene editing to generate homozygous-null lines of the two genes in two independent WT healthy control iPSC lines. Successful KO of Rab8a and Rab10 were validated by immunoblotting (Fig S1). Two clones for each Rab GTPase were further validated by Sanger sequencing and were used for subsequent experiments. Cells were differentiated into mature cortical neurons by forced expression of NGN2, according to published protocols (Sanyal et al., 2020; Zhang et al., 2013). Initially, we set out to determine whether loss of these Rab GTPases affects lysosome numbers, by LAMP1 and Lysotracker staining (Figure 1). LAMP1 staining indicated a drop in lysosome numbers in both Rab8a KO and Rab10 KO neuron models, compared to WT controls (Figure 1A). High content imaging of the Lysotracker Deep Red cell dye showed a significant decrease in Rab8a KO iNs (∼60%) in the total number of lysosomes (Figure 1B, C), while the average lysosomal area of individual lysosomes remained unaffected (Figure 1D). Lysosomal homeostasis was investigated further by assaying lysosomal pH using LysoSensor, a ratiometric pH-sensitive dye. Rab8a KO neurons exhibited significantly more acidic lysosomal lumen compared to isogenic controls (Figure 1E). In a similar line of experiments, we observed a significant decrease in lysosomal numbers in Rab10 KO compared to WT neurons, while lysosomal size was largely unaffected as in Rab8a KO neurons (Figure 1F, G, H). In contrast, we observed neutralization of lysosomal pH in Rab10 KO suggesting a divergent effect to Rab8a deficiency (Figure 1I).

**Fig 1.**
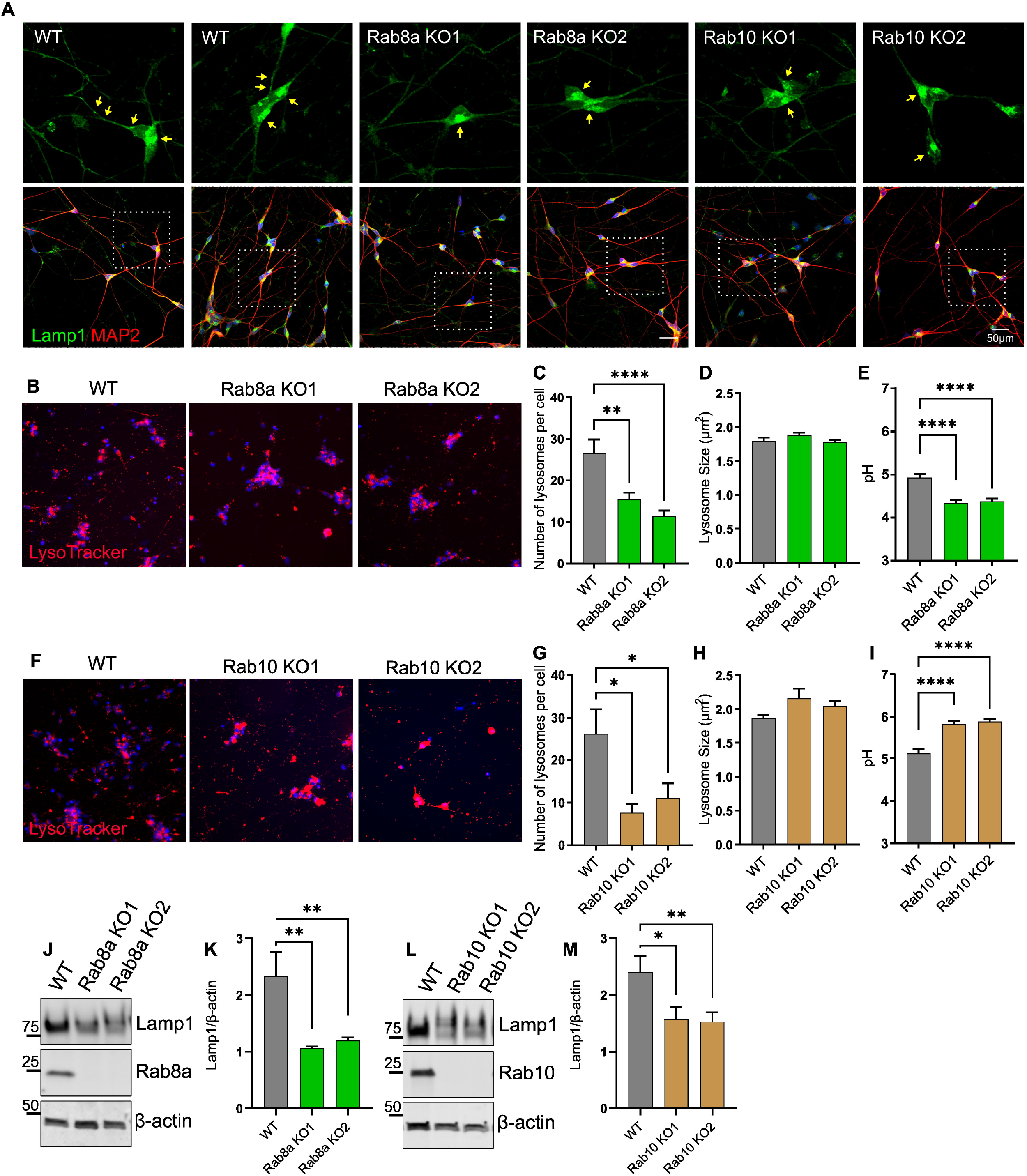
Rab8a KO and Rab10 KO human neurons have altered lysosomal morphology and function. Representative confocal images of Rab8a KO, Rab10 KO and isogenic WT control neurons stained with LAMP1, MAP2 and DAPI (A). Lysosomal parameters including lysosomal count, average lysosomal area and lysosomal pH were assessed by high-content imaging of LysoTracker Red or LysoSensor staining (B-I). Western blot analysis of levels of glycosylated LAMP1, (J-M). All lysosomal analyses were collated from 3-4 independent experiments for each of three differentiations with 10-20 wells per genotype, per experiment, on 21-day old iNs. (*, p < 0.05; **, p < 0.001; ***, p < 0.0001; ****, p < 0.0001; one-way ANOVA Tukey’s post-hoc).

As a marker of lysosomal integrity, we interrogated the levels of LAMP1 by immunoblotting. Rab8a KO and Rab10 KO neurons exhibit lower levels of LAMP1 protein running at ∼80kDa corresponding to the glycosylated protein, compared to isogenic controls (Figure 1J, K, L, M). Our data indicate a parallel effect of Rab8a and Rab10 deficiency in lysosomal numbers and levels of glycosylated LAMP1 protein but a divergent effect on lysosomal pH in human neurons. We have reported altered LAMP1 glycosylation in *in vivo* models of LRRK2 activity as well as GCase1 heterozygous-null human neurons previously (Kluss et al., 2020; Sanyal et al., 2020). Rab8a and Rab10 are involved in vesicle trafficking between the trans-Golgi network (TGN), a central organelle for protein glycosylation, and the plasma membrane. Dysregulation of LAMP1 glycosylation in our Rab GTPase KO cell models suggested impairment in TGN-related processes prompting us to assay Golgi integrity. The morphology of the CGN (GM130) and TGN (TGN46) network as well as the distribution of individual Golgi stacks and colocalization with LAMP1 were assayed by confocal microscopy. After 3D rendering of z-stack images (Imaris platform; Bitplane) CGN and TGN stacks size, relative proximity to each other and proximity to the nucleus were measured in Rab8a KO and Rab 10 KO iPSCs (Figure 2). Rab8a KO iPSCs exhibited a clustered Golgi morphology reflected in altered distribution of CGN (Figure 2A, B). Both Rab GTPase null models exhibited a decrease in total Golgi volume per cell and closer proximity to the nucleus of CGN and TGN stacks (Figure 2C-F). Our data support impairment in Golgi integrity that could affect LAMP1 glycosylation and transport to lysosomes. Analysis of LAMP1 colocalization with TGN revealed an increase in LAMP1 association with the Golgi in Rab8a KO cells but not Rab10 KO (Figure 2G) while both KO models showed a significant decrease in lysosomal numbers (Figure 2H). Taken together, these data support a model whereby inactivation of Rab8a or Rab10 is driving divergent effects on protein modification and proteostasis as well as lysosomal function that may be in part mediated by retention of LAMP1 in the Golgi.

**Fig 2.**
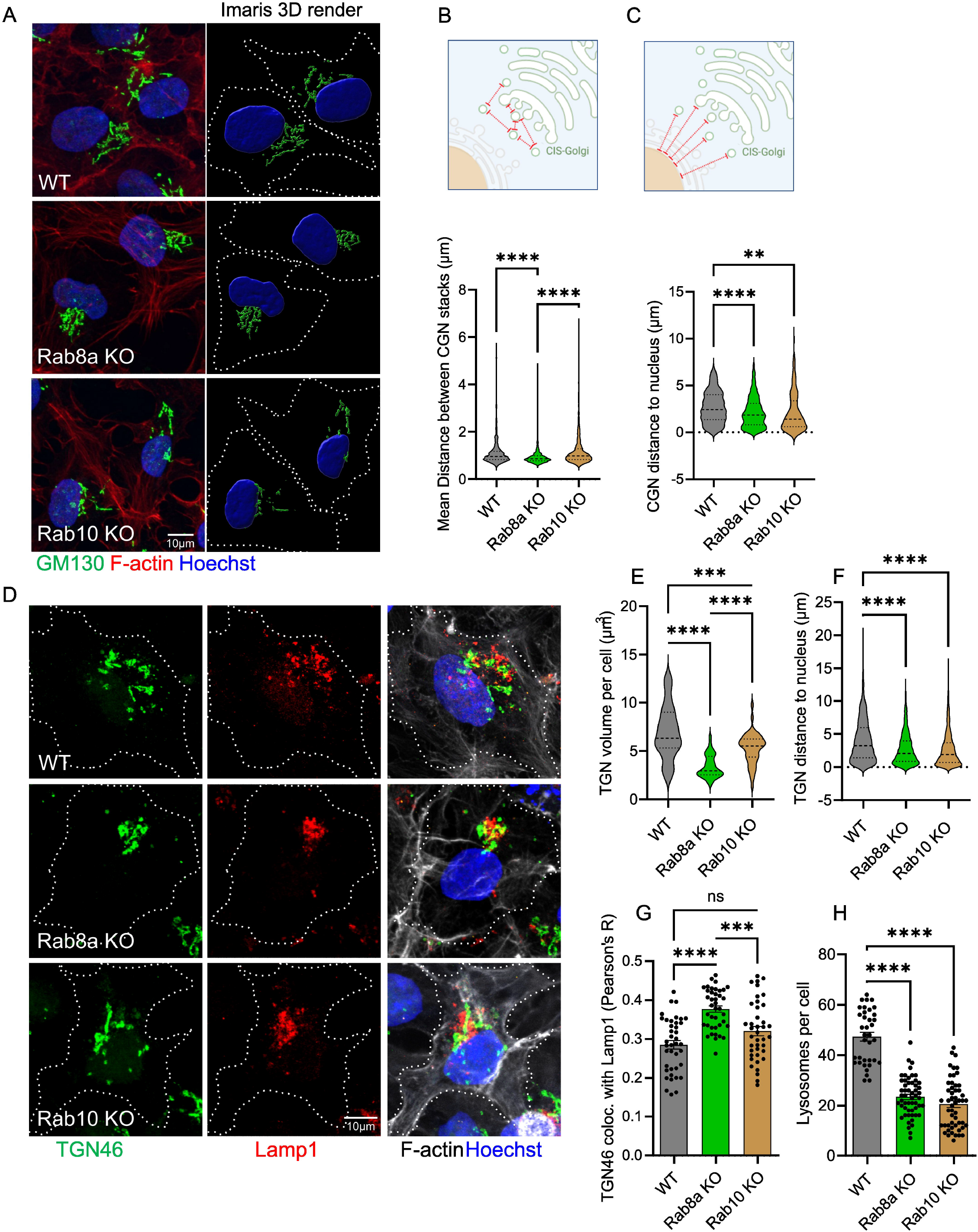
Rab8a KO and Rab10 KO cells exhibit alterations in Golgi distribution. Rab8a KO, Rab10 KO and isogenic WT control iPSCs were fixed and stained for Cis-Golgi (GM130), Trans-Golgi (TGN46) and LAMP1 (A, D). Z-stack confocal images were 3D reconstructed in Imaris (Bitplane) and the mean distance between CGN stacks (B), distance of CGN to nucleus (C), TGN volume (E), TGN distance to nucleus (F), colocalization between TGN46 and LAMP1 (G) and number of lysosomes per cell were analyzed (H). (N>50 cells over three independent experiments; **, p < 0.001; ***, p < 0.0001; ****, p < 0.0001; one-way ANOVA Tukey’s post-hoc).

### **Rab8a KO human neurons accumulate insoluble** α**-synuclein**

Accumulation of highly insoluble α-synuclein is a pathological hallmark of post-mortem PD tissue and indicative of Lewy body pathology. We, and others, have shown accumulation of α-synuclein in different neuronal models of genetic PD, including LRRK2 mutant primary neurons and *GBA1* heterozygous-null human neurons (Mazzulli et al., 2016; Sanyal et al., 2020; Schapansky et al., 2018). To assess α-synuclein metabolism in Rab8a and Rab10 null cells, we sequentially extracted total cellular proteins from iNs and determined the levels of detergent-soluble and insoluble α-synuclein. We observed an accumulation of insoluble α-synuclein in Rab8a KO neurons compared to isogenic controls while no difference was observed in Rab10 KO neurons (Figure 3A-F) suggesting Rab8a-specific effects on the capacity of these neurons to degrade α-synuclein.

**Fig 3.**
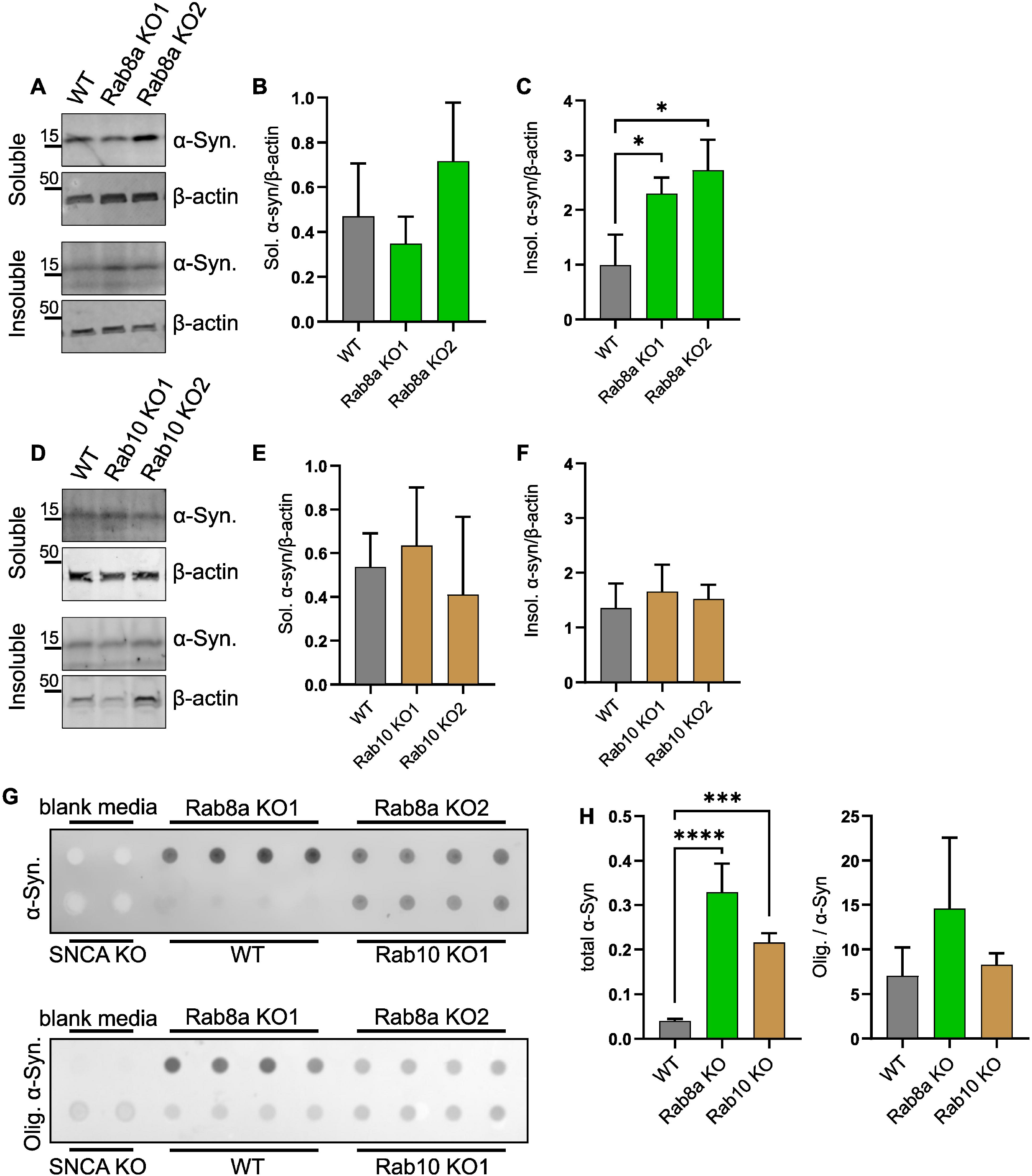
Rab8a KO human neurons accumulate insoluble α-synuclein and secrete α-synuclein in media. Rab8a KO, Rab10 KO and WT neurons were collected at D21 and extracted in NP-40 containing lysis buffer (soluble fraction), followed by SDS resuspension of the pellet (insoluble). Densitometry analysis represents α-synuclein levels normalized to actin (B, C, E, F). (G, H) Media was collected at D21 and analyzed by dot-blot for total and oligomeric α-synuclein. Protein yields of neuronal lysates were used to normalize for variability in cell density and analyze equivalent amounts of conditioned media. (*, p < 0.05; ***, p < 0.0001; ****, p < 0.0001; one-way ANOVA Tukey’s post-hoc).

One of the prominent models of how pathology may be propagated in PD supports the release of oligomeric α-synuclein from cells and direct cell-to-cell transfer (Volpicelli-Daley et al., 2014). Previously, we have shown that *GBA1* heterozygous-null neurons secrete higher levels of oligomeric α-synuclein in the media compared to isogenic controls (Sanyal et al., 2020) suggesting that lysosomal dyshomeostasis has a direct effect on α-synuclein secretion in human neurons. Here, we asked whether the increase in insoluble α-synuclein observed in Rab8a KO neurons is accompanied by changes in α-synuclein release, by immunoblotting of conditioned media. As detected by immunoblotting, Rab8a KO and Rab10 KO neurons secreted higher levels of α-synuclein in the media compared to isogenic controls, while a trend towards an increase was observed in oligomeric α-synuclein in Rab8a KO cells (Figure 3G, H). These data suggest that Rab8a and Rab10 deficiency may induce insufficient degradation and augmented secretion of synuclein species in neuronal models.

### Rab10 KO human neurons show altered Tau levels and phosphorylation

Tau pathology in the form of intracellular inclusions of hyperphosphorylated and aggregated Tau protein are a common occurrence in LRRK2 PD brain, and studies in cell models have highlighted a mechanistic interplay between Tau and Rab GTPase pathways downstream of LRRK2 activity (Guerreiro et al., 2016; Sobu et al., 2021). To characterize the effects of Rab8a and Rab10 deficiency on the biochemical properties of tau we extracted proteins from mature cortical neurons and compared the levels of Ser202/Thr205-phosphorylated Tau between the KO lines and isogenic controls. Previous studies have reported detection of distinct phospho-Tau bands, between ∼54kDa and 68kDa (Falcon et al., 2014; Jackson et al., 2016), by immunoblot analysis of human tissue. Here, we probed with a commercial pS202, pT205 Tau (AT8) antibody and detected three bands approximately at 54kDa, 62 kDa and 70 kDa (Figure 4A, B). A significant increase in phosphorylated tau (∼62kDa) was observed in Rab10 KO neurons compared to WT or Rab8a KO neurons while total tau levels were decreased (Figure 4A-D). These data demonstrate a divergence between pathways regulating disease-relevant post-translational modifications of Tau downstream of the two Rab GTPases.

**Fig 4.**
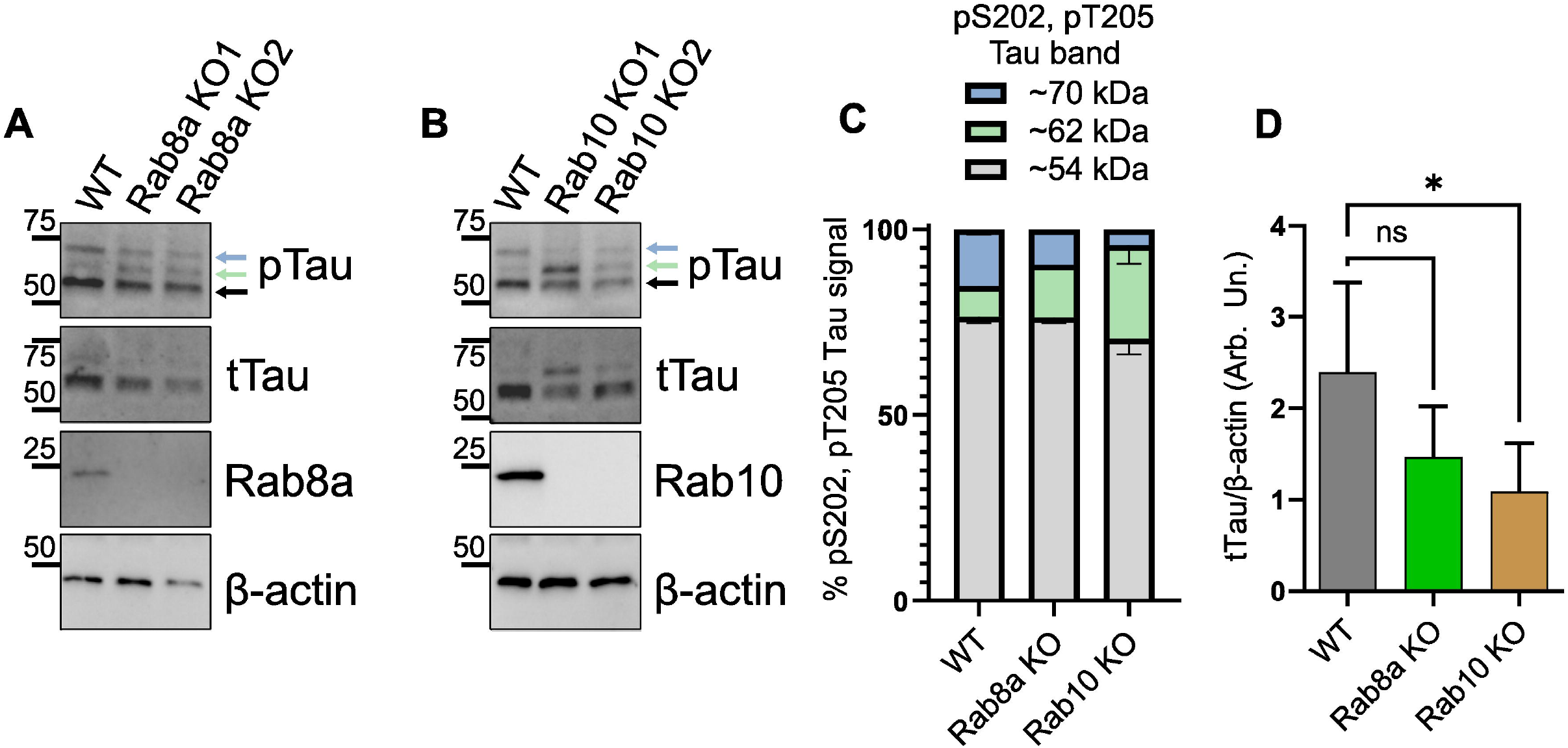
Rab10 KO increases phospho-Tau species in human neurons. Human cortical neurons were collected on D21 and phosphorylated (AT8; pTau: pS205 & pT205 Tau) and total Tau levels were assessed in Rab8a KO (A) and Rab10 KO (B) cells compared to isogenic controls (n=3 differentiations, one-way ANOVA Tukey’s post-hoc). (C) The ratio of each pTau band (54kDa, 62kDa & 70kDa) out of the total pTau signal was plotted. (D) Densitometry analysis represents total Tau levels normalized to actin. (n=3 differentiations, one-way ANOVA Tukey’s post-hoc).

## Discussion

Here we used CRISPR/Cas9 gene editing to knock out Rab8a and Rab10 in iPSCs and differentiated these cells into mature cortical neurons to examine lysosomal homeostasis and proteostasis of the pathology-relevant proteins α-synuclein and tau. LRRK2 phosphorylates Rab GTPases at a conserved residue situated in their switch II domain, and this phosphorylation is significantly enhanced by PD-associated LRRK2 mutations (Liu et al., 2018; Steger et al., 2016). Rab GTPases regulate vesicular trafficking by cycling between an inactive GDP-bound state and an active GTP-bound state, and phosphorylation by LRRK2 is believed to disrupt their interaction with GDP dissociation inhibitors thus downregulating their physiological functions (Pfeffer, 2018; Steger et al., 2016). A major limitation in studies examining signaling downstream of LRRK2 activity is that modulating LRRK2 kinase activity genetically or pharmacologically has global effects on phosphorylation and activity of >14 Rab proteins. Thus, a precise understanding of the cellular roles of individual Rab GTPases in physiological and disease-related contexts requires other strategies.

We, and others, have shown that PD-linked mutations in LRRK2 impair lysosomal homeostasis and acidification, and cause a decrease in basal autophagic flux (Schapansky et al., 2018; Mamais et al., 2021; Eguchi et al., 2018; Henry et al., 2015; Obergasteiger et al., 2020). While LRRK2 activity has been linked to the degree of lysosomal acidification, the individual LRRK2 substrates that contribute to this phenotype have not been studied. Interestingly, there is a precedent in the field for Rab8a and Rab10 to hold opposing actions. Dhekne and colleagues reported that Rab10 is a native suppressor of normal cilliogenesis upon phosphorylation by LRRK2, unlike Rab8a, and supported opposing roles for these two Rab GTPases (Dhekne et al., 2018). Here, we observed a decrease in lysosomal number per cell in both Rab8a KO and Rab10 KO neurons but a divergence in lumenal pH compared to isogenic controls. Rab8a KO cells showed higher acidification whereas Rab10 manifested alkalinization of lysosomal lumenal pH. Rab8a and Rab10 localize primarily to endosomal compartments and the TGN and regulate the dynamic interface between late Golgi membranes and endolysosomes (Goud et al., 2018; Pfeffer, 2018). LAMP1 is heavily glycosylated at the amino-terminal luminal side of the lysosomal inner leaflet, and this modification is necessary for its transport from the ER via the Golgi to the lysosome, and its function in maintaining lysosomal integrity (Schwake et al., 2013). In fact, irregular LAMP1 glycosylation has been linked to Parkinson’s and Niemann-Pick disease (Cawley et al., 2020; Fernandes et al., 2016). Here, we observed irregular glycosylation of LAMP1 in Rab8a and Rab10 KO neurons supporting impairment in LAMP1 post-Golgi trafficking. It is possible that deficiency of Rab8a or Rab10 affects trafficking of key lysosomal proteins like LAMP1 and may contribute to alterations in lysosomal pH and integrity.

Most of the terminal processing of glycoproteins is completed by glycotransferases residing in the cis-, medial- and trans-Golgi compartments and the integrity of Golgi membranes is integral to glycosylation (Stanley, 2011). Our data on irregular LAMP1 glycosylation prompted us to assess Golgi organization. Interestingly, we observed a global alteration of Golgi distribution and volume in both Rab8a KO and Rab10 KO cell models. While a previous report has shown Golgi alterations in Rab8a KO cells, Rab GTPases that are linked to membrane flux at the late Golgi/TGN-endosome interface, like Rab10, are not thought to affect Golgi organization (Aizawa and Fukuda, 2015; Pan et al., 2019). While the downstream effects of Rab8a or Rab10 deficiency on lysosomal integrity were not surprising, the parallel effect on Golgi morphology was unexpected. Importantly, we observed an increase in LAMP1 colocalization with TGN46 suggesting retention of this glycoprotein at the Golgi and impaired lysosomal biogenesis. Our observation of glycosylation and lysosomal pH defects in Rab GTPase-deficient neurons underscores the broader requirement of Rab8a and Rab10 activity in vesicular trafficking and divergent biologies downstream of LRRK2 activity.

Here, we show that levels of detergent-insoluble α-synuclein are increased in Rab8a KO neurons as well as α-synuclein release in the media in both Rab8a and Rab10 deficient human neurons. This is consistent with previous reports from our group and others showing accumulation of detergent-insoluble α-synuclein and augmented α-synuclein release in primary neurons carrying LRRK2 mutations (Schapansky et al., 2018; Volpicelli-Daley et al., 2016).

Impairment of lysosomal integrity promotes accumulation of oligomeric synuclein in cellular models highlighting the importance of the protein degradation machinery to PD pathology (Jiang et al., 2017; Lee et al., 2004). Our data suggest that LRRK2-mediated Rab8a inactivation may be sufficient to alter α-synuclein solubility.

While Lewy bodies and loss of SNc neurons are the main pathological hallmarks across PD, an overrepresentation of tau pathology has been reported in LRRK2 PD cases (Henderson et al., 2019; Herbst et al., 2022; Ujiie et al., 2012; Zimprich et al., 2004). A recent GWAS linked genetic variation in the LRRK2 locus to survival in progressive supranuclear palsy (PSP) (Jabbari et al., 2021), further highlighting a role for LRRK2 in tau pathology. Furthermore, prominent Aβ pathology is also observed in LRRK2 mutation carriers which is consistent with comorbid AD pathology (Henderson et al., 2019). How LRRK2 pathobiology might provoke distinct and independent neuropathological features is one of the greatest unanswered questions in the field. Tau is phosphorylated at multiple sites and tau aggregates associated with tauopathies are hyperphosphorylated, suggesting a role in the pathological processes that lead to aggregation (Wang and Mandelkow, 2016). While hyperphosphorylation of tau has been reported in some LRRK2 animal models (Bailey et al., 2013; Nguyen et al., 2018)the role of LRRK2 signaling on tau post-translational modification remains elusive. Here, we report an increase in Ser202/Thr205 phosphorylated tau species in Rab10 KO neurons compared to isogenic controls. Interestingly, Rab10 phosphorylation is a pathological feature in PSP and AD with reports showing colocalization between pRab10 and phosphorylated tau in neurons bearing tau-positive neurofibrillary tangles (Yan et al., 2018). In addition, Rab10 has recently been genetically linked to altered risk for AD (Andrews et al., 2019; Le Guen et al., 2021; Ridge et al., 2017; Tavana et al., 2018) further supporting an association between Rab10 pathological events and tauopathies. We observed an overall decrease in tau expression in Rab10 KO neurons compared to isogenic controls, supporting divergence in tau proteostasis. Thus, our data are in accordance with a model whereby Rab10 inactivation promotes tau phosphorylation events and may play a role in modulating tau pathology.

Our study presents novel roles for Rab8a and Rab10, two major substrates of LRRK2, and uncovers divergent phenotypes in α-synuclein and tau downstream of lysosomal dyshomeostasis. LRRK2 activity exerts an inhibitory tone across the function of multiple Rab proteins. However, when LRRK2 inhibits two Rab proteins with antagonizing features, it is more likely that one phenotype dominates while the other is suppressed. The dominant phenotype may be stochastic or more likely influenced by environmental factors and/or other genetic variables. Future work will be required to explore these phenomena across multiple pathogenic LRRK2 mutations and distinct cell types, where the outcomes may differ.

### Experimental Procedures

#### CRISPR/Cas9 Genome Editing of Human iPSCs

Single guide RNAs (sgRNAs) were selected using a web-based design tool (http://crispr.mit.edu). Rab8a sgRNA: 5’ GAACTGGATTCGCAACATTG 3’; Rab10 sgRNA: 5’ ATGGCTTAGAAACATAGATG 3’. These were cloned into pXPR_003 (Addgene #52963), modified to express the neomycin resistance gene instead of the puromycin resistance gene and sequenced using the primer 5′-GATACAAGGCTGTTAGAGAGATAATT-3′ to determine clones that successfully integrated the sgRNA. BR01 and BR33 iPSCs were generated and characterized in collaboration with the New York Stem Cell Foundation (NYSCF) using previously described methods (Muratore et al., 2017; Paull et al., 2015). BR01 and BR33 were derived from a Caucasian female and male donor respectively, who were deeply phenotyped as part of the ROS/MAP longitudinal aging studies and determined to not be cognitively impaired at death at age >89 and free from genetic variants that confer risk of PD (Bennett et al., 2018). iPSCs were co-transfected with plasmids that express dCas9 (Addgene 61425) and the sgRNA plasmid. After 2 days, cells that were successfully transfected with the two plasmids were selected by puromycin treatment for 4 days. Polyclonal cells were then monoclonally selected by plating ∼1 cell per well in a 96-well dish and allowed to grow for 2 weeks. Monoclonal lines were expanded, sequenced and stocked. Amplification of Rab8a and Rab10 genes was conducted according to manufacturer’s protocol (Invitrogen K2030-01) and homozygous gene editing was confirmed by Sanger sequencing. Multiple sequence alignment was performed using ClustalW in BioEdit.

#### Differentiation of Human iPSCs

iPSCs were cultured in StemFlex (A33493) media. 100,000 cells/cm3 were co-transduced with lentivirus packaged with pTet-O-NGN2-puro and Fudelta GW-rtTA plasmids (Zhang et al., 2013) for 2 days and passaged for expansion/stocks. NGN2-transduced iPSC cells were thawed in StemFlex media with ROCK inhibitor (10 μM; Stemcell Technologies, 72304) and plated at 2 × 10^6^ cells/10 cm plate and grown until 75% confluent. For differentiation, on day 1 cells were fed with KnockOut media (Gibco 10829.018) supplemented with KnockOut Serum Replacement (Invitrogen 10928-028), 1% MEM non-essential amino acids (Invitrogen 11140), 1% GlutaMAX (Gibco 35050061) and 0.1% BME (Invitrogen 21985-023) (KSR) with doxycycline (2 μg/ml, Sigma, D9891-5g) to induce NGN2 expression. On day 2, they were fed with a 1:1 ratio of KSR:N2B media (DMEM F12 supplemented with 1% GlutaMAX, 3% dextrose and N2-Supplement B; StemCell Technologies 07156) with puromycin (5 μg/ml; Life Technologies, A11138-03) and doxycycline to select for transduced cells. On day 3, the cells were fed with N2B media with B27 (1:100; Life technologies, 17504-044), puromycin, and doxycycline. On day 4, induced neurons (iNs) were frozen down in 10% DMSO/FBS in Neurobasal media (NBM Gibco 21103-049) supplemented with B27, BDNF (Peprotech, 450-02), CNTF (Peprotech, 450-13), and GDNF (Peprotech, 450-10) all at 100 ng/uL, ROCK inhibitor (10 μM), puromycin, and doxycycline. iNs were plated and grown in NBM with B27, BDNF, CNTF, GDNF, puromycin, and doxycycline until day 21.

#### Western Blot

Cells were lysed in cell lysis buffer (50 mM Tris-HCl, 150 mM NaCl, 0.5 mM EDTA, 0.5% (v/v) sodium deoxycholate, 1% (v/v) NP-40, pH 8) with protease and phosphatase inhibitors for 30 min. Lysates were centrifuged at 14,000g for 15 min at 4°C, and supernatants were quantified using the BCA assay. Samples were prepared in 1x SDS-PAGE loading buffer, denatured at 65°C for 10 min, resolved on an SDS-polyacrylamide gel, and transferred to PVDF membrane. Membranes were blocked with 5% BSA (Sigma) prior to probing with primary antibodies against LAMP1 (abcam ab108597), Rab10 (abcam ab237703), Rab8a (Cell Signalling 6975), αSyn oligomer (abcam ab209538), total αSyn (MABN1817), pTau (Ser202, Thr205) (Thermo; MN1020), and total Tau (13-6400). Secondary antibodies conjugated to horseradish peroxidase were used for detection using autoradiography.

#### Immunocytochemistry

Neurons or iPSCs were seeded at 120,000 cells/well (24 well plate) on 12 mm coverslips precoated with Geltrex basement membrane matrix (A1413201, Thermo) and cultured as described above. Cells were fixed in 4% (w/v) formaldehyde/PBS for 15 minutes, permeabilized in 0.2% Triton X-100/PBS for 10 minutes at RT, blocked in 5% (v/v) FBS in PBS, and incubated with primary antibodies in 1% (v/v) FBS/PBS overnight at 4C. Following 3 washes in PBS, the cells were incubated for 1 hour with secondary antibodies (Alexa Fluor 488, 568, 647-conjugated; ThermoFisher). After 3 PBS washes, the coverslips were mounted, and the cells were imaged by confocal microscopy (Olympus Fluoview FV-1000). The antibodies used were TGN46 (AHP500; BioRad), Lamp1 (DSHB; H4A3), GM130 (Cell Sign. Tech.; 12480). The

Coloc2 ImageJ plugin was used for analysis of the Pearons overlap coefficient r. For high-content imaging, neurons were plated 10,000 cells per well in 96-well black-wall clear-bottom plates (Greiner), labeled with LysoTracker Red (Invitrogen) according to the manufacturer’s specifications, and 20 ng/ml of Hoechst. Labeled live cells were imaged at 10x magnification, six fields per well, in the DAPI and Cy3 channels using an IN Cell Analyzer 2200 (GE Healthcare). Images were analyzed with the IN Cell Workstation software (GE Healthcare) multi target analysis protocol. Briefly, nuclei were segmented by applying a Top Hat algorithm with a minimum area of 50 square μm and a sensitivity level of 50 to the DAPI channel. Lysosomes were defined as objects with a 1–3 μm diameter, segmented by 2 scales with a sensitivity level of 20 in the corresponding channel. Cell count, lysosome count, mean lysosome area and total lysosome area were calculated.

#### Imaris analysis

Following confocal microscopy, the Imaris platform (Bitplane, Zürich, Switzerland) was used to analyze the localization and distribution of CGN, TGN and LAMP1. In Fig 2, z-stack confocal images of iPSCs were processed through the Imaris Surface Contour module to render Golgi, LAMP1 and nuclear staining to surfaces and measure the distance between CGN/TGN stacks and proximity to the nuclear edge throughout z planes in the 3D volume.

#### LysoSensor Assay

For lysosomal pH analysis, the ratiometric dye LysoSensor Yellow/Blue (Invitrogen) was used. Neurons were plated at 10,000 cells per well on 96-well black-wall black-bottom plates (Thermo Scientific), labeled with dye (1 μM) for 10 min prior to rinsing 2x with HBSS buffer. Cells were imaged using a Synergy H1 hybrid reader (Bio-Tek; reading at excitation 329/384, emission 440/540). Then, cells were incubated for 5 min at 37°C with pH calibration standards (pH of 3.96, 4.46, 4.96, 5.47, and 5.97) prepared in 20 mM 2-(N-morpholino)ethanesulfonic acid, 110 mM KCl, and 20 mM NaCl freshly supplemented with 30 μM nigericin and 15 μM monensin. A pH standard curve was determined for each genotype using GraphPad Prism 7 and individual baseline pH values were interpolated from these standard curves.

#### Dot-blot assay

Media was collected from cells at D21 after being conditioned for 48 hours, centrifuged at 500g for 5 mins and amounts normalized to cell density/protein load were diluted into 300μL TBS and added to the respective wells on 0.2μm nitrocellulose (Bio-rad, 1620112). The membrane was dried for 10 mins at RT before fixation in 2% PFA for 30 mins, blocking in 5% BSA and probing with primary antibodies overnight at 4°C.

#### Statistical Analyses

All experiments were conducted at least three independent times for three differentiations. Error bars indicate mean +SD. Statistical analysis was performed using GraphPad Prism software, using a one-way ANOVA with Tukey’s post-hoc test.

## Figure legends

**Supplementary Fig 1. Validation of Rab8a KO and Rab10 KO clones by Western blot and Sanger sequencing**. Validation of successful KO of Rab8a (A) and Rab10 (B) by Western blot. Two clones of each Rab GTPase edited line were selected for further validation by Sanger sequencing and experimentation (C).

## Acknowledgments

The authors would like to thank Dr. Matthew Farrer (UF/CTRND) for access to the Olympus Fluoview FV-1000 confocal laser scanning microscope used for imaging.

## Author contributions

MJL, AS, AM: Conceptualization, Methodology, Data Analysis, Writing-Original draft preparation. AM, AS, AF, CGZ, MG, LRD, NS, WG, SL: Visualization, Investigation. AM, MJL: Supervision. AF, AM, MJL: Writing-Reviewing and Editing

