## Supplementary figures and images for "The LRRK2 kinase substrates Rab8a and Rab10 contribute complementary but distinct disease-relevant phenotypes in human neurons"

### Supplementary Figure 1

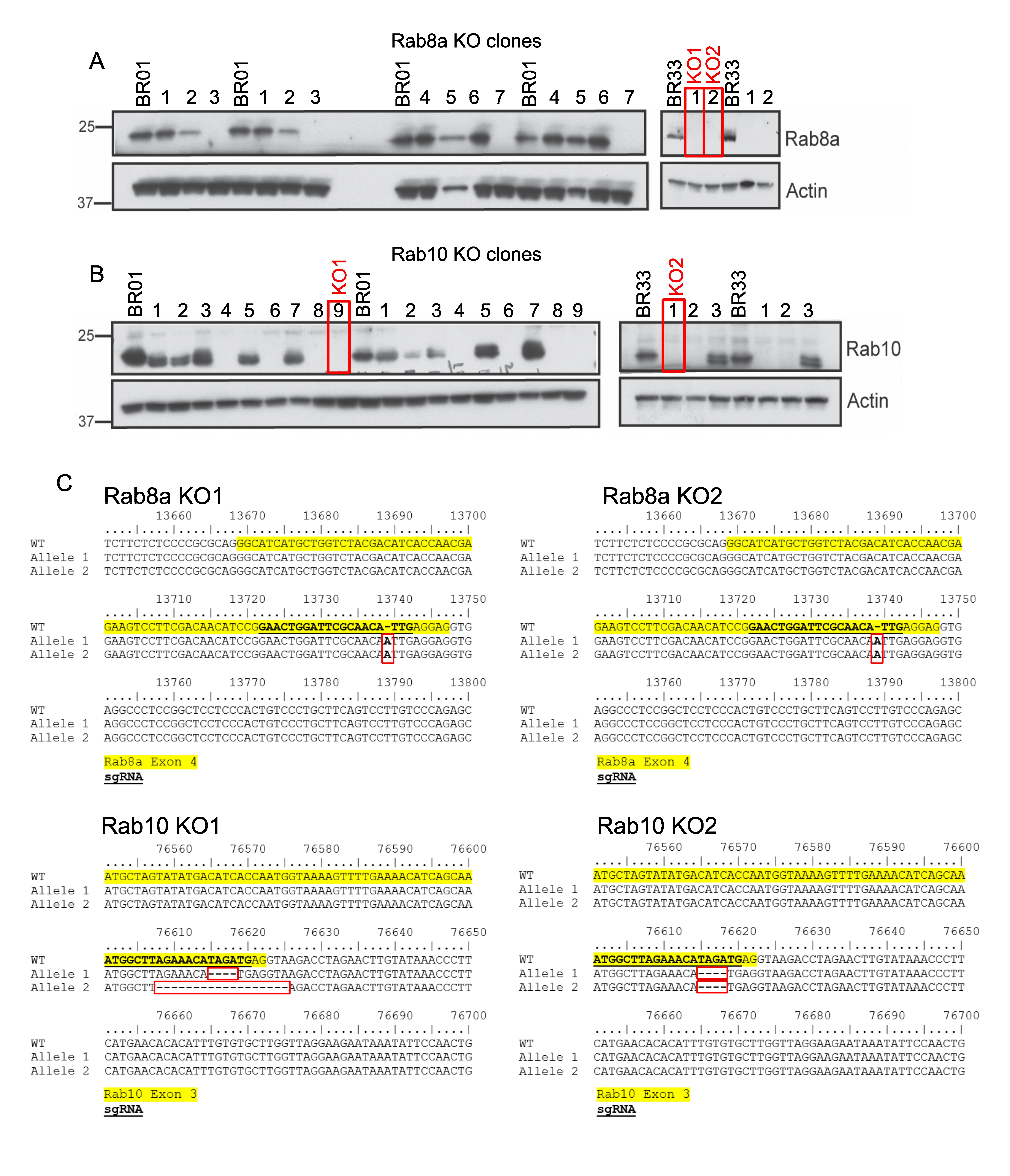
